# Transport and Organization of Individual Vimentin Filaments Within Dense Networks Revealed by Single Particle Tracking and 3D FIB-SEM

**DOI:** 10.1101/2024.06.10.598346

**Authors:** Bhuvanasundar Renganathan, Andrew Moore, Wei-Hong Yeo, Alyson Petruncio, David Ackerman, Aubrey Wiegel, The CellMap Team, H. Amalia Pasolli, C. Shan Xu, Gleb Shtengel, Harald F. Hess, Anna S Serpinskaya, Hao F. Zhang, Jennifer Lippincott-Schwartz, Vladimir I. Gelfand

## Abstract

Vimentin intermediate filaments (VIFs) form complex, tight-packed networks; due to this density, traditional ensemble labeling and imaging approaches cannot accurately discern single filament behavior. To address this, we introduce a sparse vimentin-SunTag labeling strategy to unambiguously visualize individual filament dynamics. This technique confirmed known long-range dynein and kinesin transport of peripheral VIFs and uncovered extensive bidirectional VIF motion within the perinuclear vimentin network, a region we had thought too densely bundled to permit such motility. To examine the nanoscale organization of perinuclear vimentin, we acquired high-resolution electron microscopy volumes of a vitreously frozen cell and reconstructed VIFs and microtubules within a ∼50 µm^3^ window. Of 583 VIFs identified, most were integrated into long, semi-coherent bundles that fluctuated in width and filament packing density. Unexpectedly, VIFs displayed minimal local co-alignment with microtubules, save for sporadic cross-over sites that we predict facilitate cytoskeletal crosstalk. Overall, this work demonstrates single VIF dynamics and organization in the cellular milieu for the first time

**Summary:** Single-particle tracking demonstrates that individual filaments in bundles of vimentin intermediate filaments are transported in the cytoplasm by motor proteins along microtubules. Furthermore, using 3D FIB-SEM the authors showed that vimentin filament bundles are loosely packed and co-aligned with microtubules.

## Introduction

Intermediate filaments (IFs) are a diverse class of cytoskeletal polymer encoded by a family of over 70 distinct genes (Szeverenyi et al., 2008). So named due to their intermediate width relative to actin filaments and microtubules, IF proteins assemble long nonpolar fibers with widths of ∼10-12 nm (Steinert et al., 1985; Eibauer et al., 2024). Type III IFs comprise a closely related subset of these proteins, which form mature assemblies within the cytoplasm and display striking cell type specificity (Herrmann et al., 2007; Redmond and Coulombe, 2021; Omary et al., 2004). Vimentin, perhaps the best studied type III IF proteins, is canonically expressed in mesenchymal cells such as fibroblasts, and forms complex, multifunctional networks (Lowery et al., 2015).

Due to their abundance, unique distribution, and biophysical properties, Vimentin IFs (VIFs) are ideally suited to organize cytoplasmic space. VIFs interact with actin and microtubules, fostering cytoskeletal crosstalk required to adapt cell shape and organelle distribution to ongoing cellular needs (Lynch et al., 2013; Kim and Coulombe, 2007). VIFs also interact with an ever-growing list of membranous organelles, including the Golgi apparatus (Vitali et al., 2023), mitochondria (Nekrasova et al., 2011), and the nuclear envelope (Patteson et al., 2019), to modulate organelle structure, dynamics, and subcellular positioning (Lowery et al., 2015; Chang et al., 2009). Accordingly, pathological conditions that interfere with VIF structure or dynamics are often marked by disordered subcellular organization (Renganathan et al., 2023).

VIF distribution within the cytoplasm is notably non-uniform, exhibiting a radial gradient of filament density. In the perinuclear region, dense, interconnected VIF bundles crisscross the cytoplasm, while at the cell periphery sparser and simpler filament arrangements predominate. When first characterized by early immunocytochemistry studies, this unique organization of VIF networks was presumed to reflect a largely static immobile character (Starger et al., 1978; Franke et al., 1978). This interpretation was revised upon the development of live fluorescence microscopy techniques that allowed for timelapse movies of fluorescent protein tagged vimentin in living cells (Ho et al., 1998, Yoon et al., 1998, Prahlad et al., 1998). Live imaging and the use of techniques such as fluorescence recovery after photobleaching (FRAP) and targeted photoactivation/photoconversion, demonstrated that vimentin filaments move along microtubules. Vimentin motility was subsequently shown to play key roles in cell migration and wound healing (Eckes et al., 2000; Cheng et al., 2016; Gan et al., 2016)

Though these techniques were invaluable in identifying and defining the ensemble behavior of the vimentin cytoskeleton, they are limited in their ability to report on the motility of single filaments within the broader VIF network. As single filaments are ∼10-12nm in diameter, it is impossible in conventional light microscopy movies to determine whether elongated, threadlike vimentin structures are true single filaments or bundled arrays. In the perinuclear region, where vimentin filament density is highest, the problem is further compounded. To investigate the behavior of single vimentin filaments the field has largely favored in vitro reconstitution assays, in which VIFs are assembled in a cell free system and assayed by Total Internal Reflected Fluorescence (TIRF) microscopy (Winheim et al., 2011). These assays have shed light on fundamental VIF properties helping to define the fundamental biomechanical profile of these polymers. However, single VIF dynamics are underexplored in their native environment, intact cells.

In this study we used single particle tracking and volume EM reconstruction to define the motion and organization of individual vimentin filaments in cultured cells. First, we developed a dual labeling strategy to visualize ensemble vimentin organization using a standard vimentin-mCherry reporter whilst simultaneously tracking single VIF motion via an ultrabright vimentin-SunTag fusion. By tuning vimentin-SunTag expression level through sparse labeling and short transfection windows, we achieved labeling densities at or below a single reporter per filament. We simultaneously tracked hundreds of single filaments across the cell, including in the hyperdense perinuclear region that has been historically challenging to study. VIF motility was abrogated by chemical depolymerization of microtubules or inhibition of either dynein or kinesin-1 microtubule motors. To our surprise, we observed clear bidirectional motion of single filaments within dense bundles, suggesting that VIF bundles are not tightly cross-linked. To further explore the structural organization of dense, perinuclear vimentin we turned to isotropic volume EM of a cryofixed, freeze substituted cells. With sufficient contrast to identify and track vimentin filaments, this approach provided a window into the precise organization of intermediate filaments and the microtubules surrounding them in an intact cell.

## Results

### Vimentin-SunTag labels the mature VIF

To explore the dynamics of single vimentin filaments we developed a simultaneous, two-color labeling strategy in RPE cells. Transfected vimentin-mCherry co-assembles with endogenous VIFs and is a well-established reporter of vimentin distribution. In parallel, we co-transfected cells with our single molecule reporter, a fusion between human vimentin and SuperNova tag (SunTag), a synthetic scaffold harboring 24 GCN4 peptide epitope (Tanenbaum et al., 2014). Vimentin-SunTag is expressed under the control of a weak CMV promotor and co-expressed with an additional construct encoding anti-GCN4 single chain fragment variable (scFV) fused to superfolder-GFP (sfGFP) and a nuclear localization signal to limit excess cytosolic background. Once translated, the scFV-GCN4-sfGFP associates with the arrayed GCN4 epitopes, massively amplifying the fluorescence signal associated with a single vimentin monomer. By tuning expression of vimentin-SunTag we could achieve remarkably low expression levels of the probe with 4.35 ± 0.6 SunTag spots per square micron (mean +/-sd) a density consistent with one or fewer SunTag-labeled vimentin monomers per filament.

Using this approach, we were able to identify and track single vimentin-SunTag labeled filaments and contextualize their dynamics within their complex, highly overlapping vimentin-mCherry labeled neighborhood (Fig. 1A). The SunTag system is particularly well suited for tracking single vimentin filaments, compared to single dyes or photoactivatable proteins, as its multimeric nature allows for reconstruction of substantially longer single particle trajectories than would be possible with a single fluorophore (Fig. 1B and Video 1).

**Figure 1:**
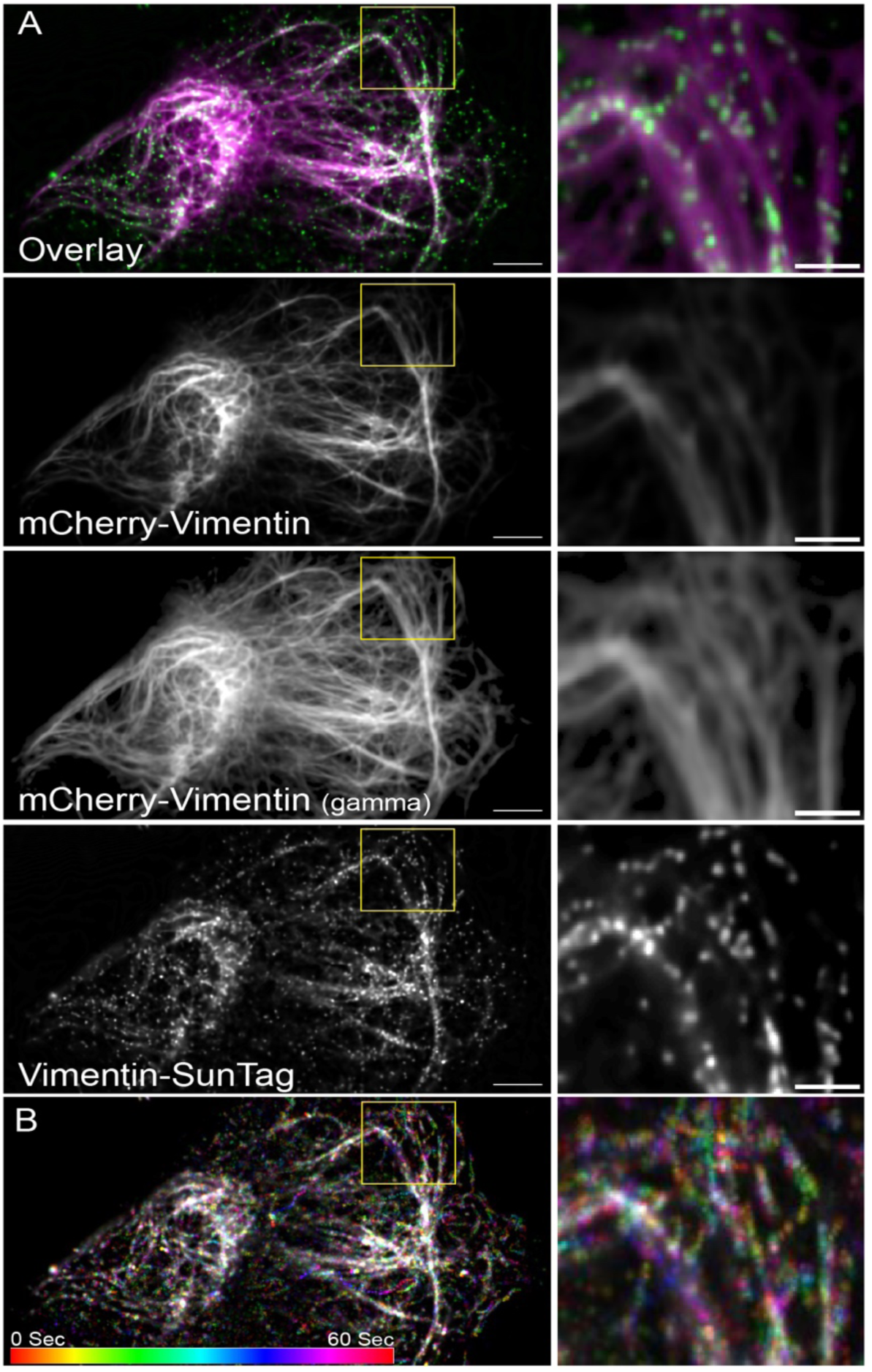
Vimentin-SunTag incorporates into endogenous vimentin filaments. Cells co-transfected with Vimentin-SunTag constructs and vimentin-mCherry plasmid were imaged 24 hours post-transfection. Movies were collected for 1 minute at 2 frames per second (fps). A) Snapshot of the first frame (0 sec) from Video 1. Endogenous vimentin organization is visualized with vimentin-mCherry in magenta. Vimentin-SunTag in green decorates the length of endogenous vimentin filaments. The yellow box is zoomed in and shown in the right panel. Gamma correction was applied to vimentin-mCherry to visualize the thin vimentin filaments. B) Pseudo-colored maximum intensity time projection of the 60-second, 120-frame Vimentin-SunTag movie with an accompanying color bar. The yellow box is zoomed in and shown in the right panel. Scale bar: 5 µm for A and 1 µm for insets.

### Vimentin-SunTag incorporates into mature VIF

To validate this approach and demonstrate that our reporter accurately integrates into VIFs, we fixed cells expressing vimentin-SunTag and performed immunostaining to visualize endogenous vimentin. In structured illumination microscopy (SIM) images we observed vimentin-SunTag puncta distribution throughout the cytoplasm and in closely alignment with the position of endogenous VIFs (Fig. 2A). The integration of vimentin-SunTag into endogenous VIFs was further confirmed using platinum replica electron microscopy (PREM). Cells expressing vimentin-SunTag were extracted with Triton X-100, chemically fixed and stained using indirect immunofluorescence with an anti-GFP primary antibody and 12nm colloidal gold conjugated secondary antibody. Direct microscopic observation shows that Triton extraction did not remove any SunTag particles from the cells. Cells were then prepped for PREM including critical point drying and rotary shadowing (see methods for details), and transmission EM images were collected. Control cells that were not transfected with vimentin-SunTag exhibited no gold particles (Fig. 2B), while vimentin-SunTag-transfected cells displayed clear 12 nm electron-dense particles associated with individual VIFs, consistent with the size of the colloidal gold particles (Fig. 2C). VIF organization appeared comparable between control and experimental groups, indicating that the structure of endogenous VIFs was not disturbed by our reporter’s expression. Together our SIM and PREM data provide clear evidence that vimentin-SunTag integrates into endogenous VIFs and does not inhibit normal filament assembly. Having validated that our probe was indeed localized to single VIFs, we next sought to track the motile behavior of SunTag spots to analyze the motility pattern of individual vimentin filaments.

**Figure 2:**
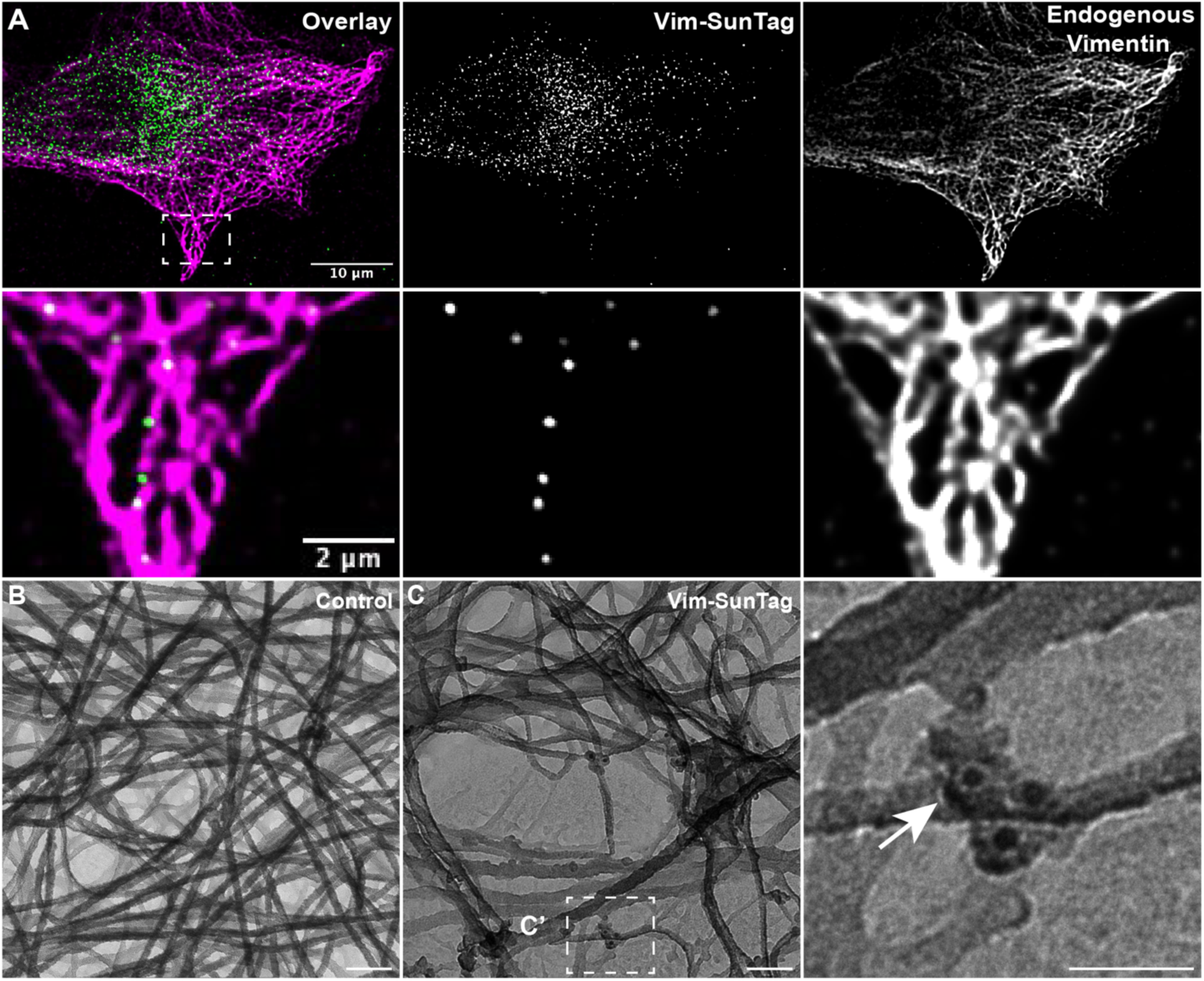
Vimentin-SunTag incorporates into mature vimentin filaments. (A) Structured Illumination Microscopy (SIM) micrograph of a fixed RPE cell expressing Vimentin-SunTag (green) and immunolabeled for endogenous vimentin (magenta). The enlarged inset highlights colocalization of vimentin-SunTag puncta with endogenous vimentin. Platinum replica electron microscopy (PREM) micrograph of a (B) control cell and (C) vimentin-SunTag expressing cell. Black dots in panel C indicate immunogold particles tagged with anti-GFP antibodies. The white dotted box in panel C was enlarged and shown as C’. The white arrow points to immunogold particles decorating the endogenous vimentin filaments. Immunogold labeling clearly demonstrates that expressed vimentin-SunTag incorporates into individual vimentin filaments. Scale bar: 100 nm for E & F; 50 nm for F’-F”. Scale bar 10 µm (A), 2 µm (A inset), 100 nm (B and C), 50 nm (C insets).

### Vimentin-SunTag Reveals Robust Transport of VIF

To analyze motion of single VIFs, we transfected RPE cells with vimentin-SunTag, and collected timelapse movies at 2 frames per second (fps) over 1-minute spans. We hypothesized that vimentin-SunTag would be highly mobile in the cell periphery, but largely immobilized in the dense, perinuclear vimentin cytoskeleton. To our surprise, we observed robust vimentin-SunTag motility in all regions of the cell (Video 1 and 2), suggesting a much greater level of baseline VIF motion than we had anticipated (Fig. 3A and B). Vimentin-SunTag puncta displayed both anterograde and retrograde movement (Fig. 3C).

**Figure 3:**
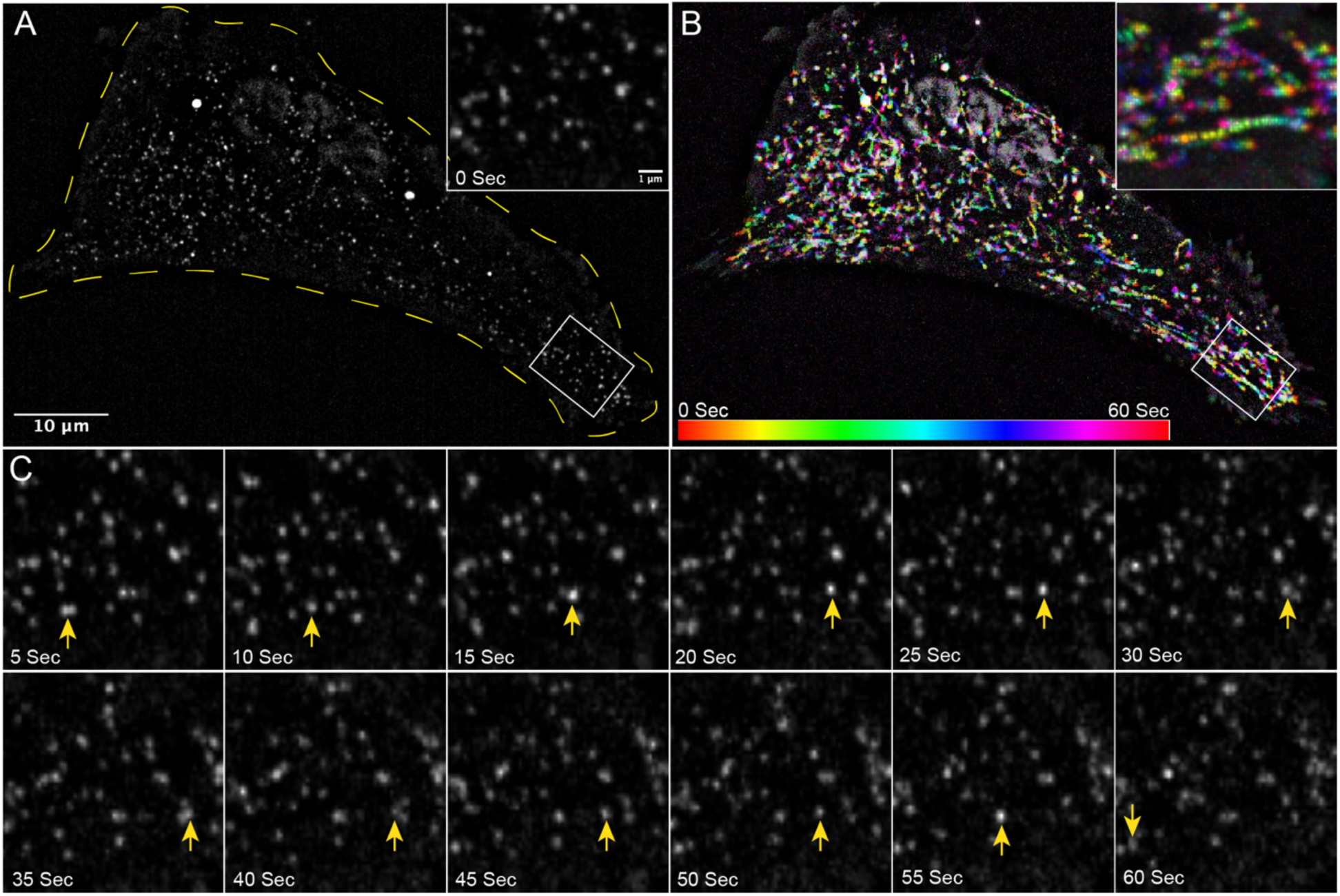
Vimentin filaments are highly motile. Vimentin-SunTag constructs were transfected into RPE cells, 24 hours post-transfection, movies were collected for a total time of 1 minute at 2 frames per second (fps) (See also Video S1). (A) Snapshot of first frame (0 sec) from Video S1, displays the distribution of vimentin-SunTag (0 sec). The yellow line indicates cell boundary. White box has been zoomed and shown as inset. (B) Pseudo-colored maximum intensity time projection of 60 second, 120 frame vimentin-SunTag movie with accompanying color bar. White box has been zoomed and shown as inset. (C) Montage of the zoomed region from panel A, displaying snapshots at five-second intervals. A yellow arrow indicates the position of a specific vimentin-SunTag dot across different snapshots of the movie. Scale bar: 10 µm for A and B; 1 µm for inset in A, B and for snapshots in panel C.

Single vimentin-SunTag dots were tracked, and their track length was quantified using particle analysis software. Based on the track length over the total imaging time (1 min), we classified them into three categories: long linear trajectories (>1.0 µm), short displacements (0.3 to 1.0 µm), and stationary dots (<0.3 µm). Among the total vimentin-SunTag dots analyzed, 13% exhibited long trajectories, 64% displayed short displacements, and 22% were identified as stationary dots. Notably, during migration along long trajectories, vimentin-SunTag occasionally paused or underwent a reversal of direction before continuing (Fig. 3C, 4A and B, video 1 and 2).

### VIF transport depends on microtubules

We have previously demonstrated that global VIF transport is dependent on microtubule tracks (Hookway et al., 2015). Given this dependence, we aimed to determine if individual filament motion detected by our Vimentin-SunTag reporter similarly requires microtubules. To explore this, we employed nocodazole, a microtubule depolymerizing agent, to acutely disrupt the microtubule network. Chronic, long-term microtubule depolymerization is known to eventually cause vimentin cytoskeleton collapse into the perinuclear region (Goldman 1971); however, Gan et al. (2016) demonstrated that normal VIF distribution can persist for at least 30 minutes following nocodazole treatment. Consistent with this, treatment of RPE cells with 10 µM nocodazole depolymerized most microtubules within 15 minutes, yet the VIF network remained mostly intact and did not collapse for up to 30 minutes (Fig S1). Using our Vimentin-SunTag reporter, we then queried whether VIF movement would be visible during this 15-minute time window between microtubule loss and VIF collapse, which would suggest non-microtubule-based motion. Vimentin-SunTag exhibited restricted jiggling, but long trajectories were almost rarely observed (Fig. 4 C and D, Fig. S2 A, and Video 3). Quantification revealed a drastic 4.5-fold decrease in long linear tracks and a 2.8-fold increase in stationary dots without microtubules compared to the control condition (Fig. S2 E).

**Figure 4:**
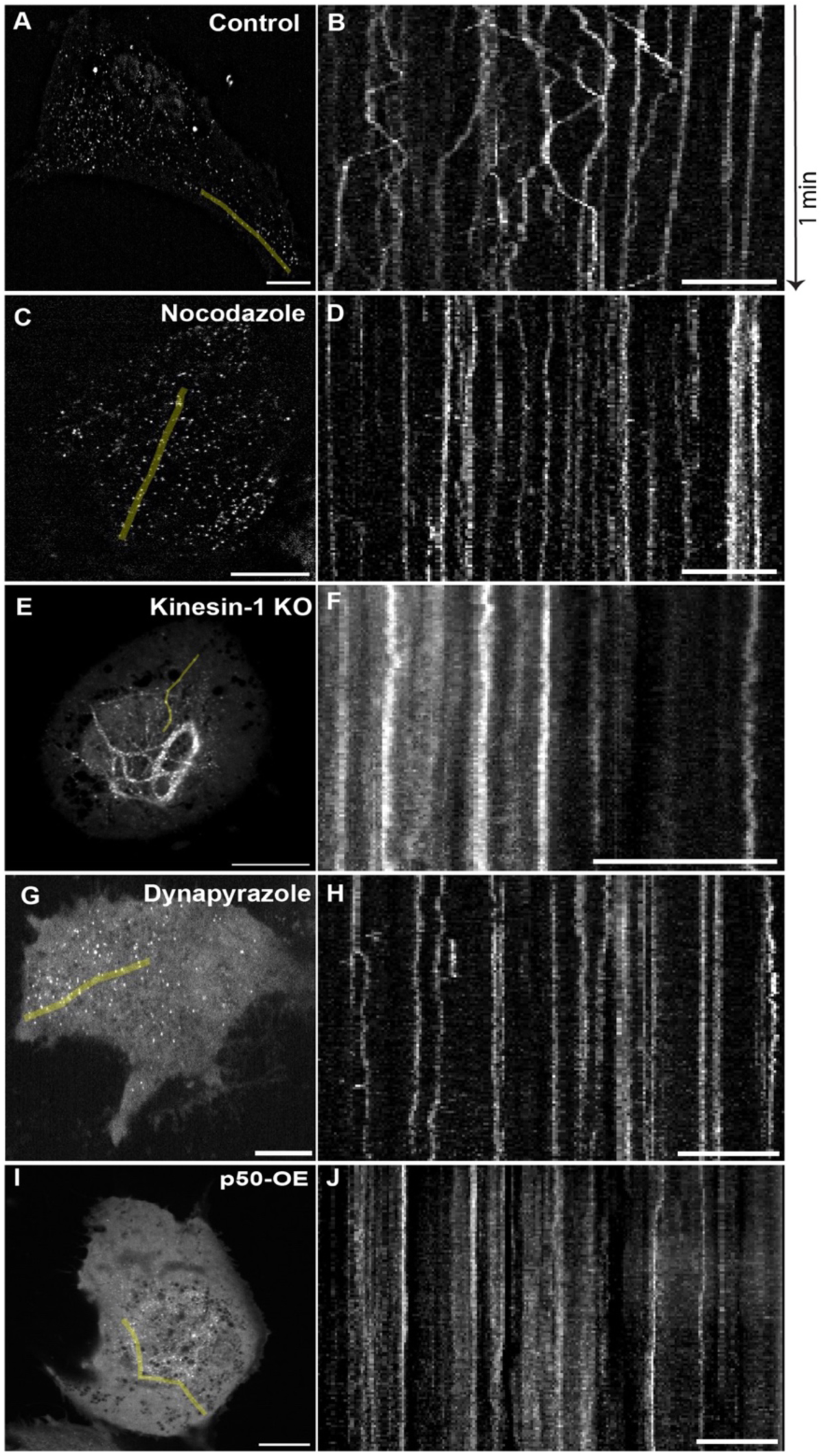
Microtubule Motors Kinesin-1 and Cytoplasmic Dynein Drive Bidirectional Vimentin Filament Transport. RPE cells were transfected with vimentin-SunTag. After twenty-four hours, a time-lapse movie was captured at 2 fps for a total duration of 1 minute. The region of kymograph generated from each time sequence is represented as green line. Scale bar in microscopic images (A, C, E, G, and I) represents 10 µm and in kymographs (B, D, F, H, and J) represents 5 µm. A) First frame of the image sequence from a control cell (Video S2), and B) Kymograph from the control. C) First frame of the image sequence from a cell treated with 10 µM of Nocodazole (Video S3), and D) Kymograph from the Nocodazole-treated cell. E) First frame of the image sequence from a kinesin-1 knockout (KO) cell (Video S4), and F) Kymograph from the kinesin-1 KO cell. G) First frame of the image sequence from a cell treated with 5 µM of Dynpyrazole A (Video S5), and H) Kymograph from the Dynapyrazole A treated cell. I) First frame of the image sequence from a p50 overexpressing (OE) cell (Video S6), and J) Kymograph from the p50 overexpressing cell.

### Microtubule motors kinesin-1 and cytoplasmic dynein drive VIF transport

The proper distribution of VIFs along microtubules is dependent on kinesin-1 (Gyoeva and Gelfand, 1991). In RPE cells, the KIF5B isoform of kinesin-1 is predominantly expressed, and its knockout inhibits VIF transport, leading to their perinuclear accumulation, even in the presence of intact microtubules (Robert et al., 2019). Here we tested the behavior of our vimentin-SunTag reporter in RPE KIF5B null cells. Consistent with our earlier results (Robert et al., 2019), most SunTag dots were clustered in the perinuclear region (Fig. 4 E and F, Fig. S2 B, and Video 4), with a 4-fold increase in stationary dots and a 3.2-fold decrease in long-linear tracks compared to the control condition (Fig. S2 E).

We reasoned that cytoplasmic dynein, the primary retrograde microtubule motor, is likely responsible for the long-range transport of SunTag-labeled single vimentin filaments towards the nucleus. To investigate this hypothesis, we visualized the dynamics of single VIFs upon either chemical or genetic inhibition of dynein. Acute dynein inhibition with 5 µM Dynapyrazole A for 45 mins eliminated all long-range vimentin-SunTag trajectories, significantly increasing the fraction of spots displaying a confined motility pattern (Fig. 3 G and H, Figs. S2 C, and Video 5). As a complementary strategy, we overexpressed the p50 component of the dynactin complex to inhibit cytoplasmic dynein (Burkhardt et al., 1997) and observed comparable deficits in long-range transport (Fig. 3 I and J, Figs. S2 D, and Video 6). Compared to control cells, inhibition of dynein by either approach resulted in a decrease in long tracks by 4.8- and 3.9-fold, respectively, as well as an increase in confined, stationary spots by 3.3- and 2.7-fold, respectively (Figs. S2 E). Thus, the inhibition of either of the two major microtubule motors, kinesin-1 and cytoplasmic dynein, results in the impairment of long-range VIF transport. Notably, dynein inhibition by p50 overexpression resulted in perinuclear vimentin accumulation, reminiscent of the pattern we observed in KIF5B-null cells. That inhibition of a retrograde motor should induce a perinuclear accumulation phenotype was unexpected, and strongly suggested that the activities of kinesin and dynein in VIF transport are tightly coupled.

### Nanoscale three-dimensional reconstruction of VIF networks in intact cells

The bidirectional trafficking of individual filaments within dense regions of the vimentin cytoskeleton surprised us, as we had anticipated these areas would consist primarily of highly compact bundles composed of ordered arrays of closely apposed vimentin filaments. We reasoned that because the motility pattern observed with our reporter challenged our preconceived model of VIF bundle organization, we needed to critically reexamine the nanoscale structure of vimentin filaments in their native cellular context. Recognizing that even super resolution light microscopy would not allow us to fully appreciate the fine organization of vimentin filament bundles (Fig. S3 A), we turned to Focused Ion Beam Scanning Electron Microscopy (FIB-SEM). Cells were prepared as in Xu et al. 2021. In brief, cells were cultured on sapphire coverslips and cryofixed by high-pressure freezing. Frozen cells were subsequently freeze-substituted in media containing osmium tetroxide and uranyl acetate before final mounting in Durcupan. FIB-SEM imaging was performed over seven days with xy pixel size set to 2 nm and focused Ga+ beam set to ablate 2nm of the sample surface per cycle. The raw stack was then aligned, registered, and scaled, yielding an isotropic 2 x 2 x 2 nm voxel stack, from which we selected a 5.0 x 3.0 x 3.6 µm volume in the perinuclear region (Fig. 5 A) for in-depth filament tracing and analysis.

**Figure 5:**
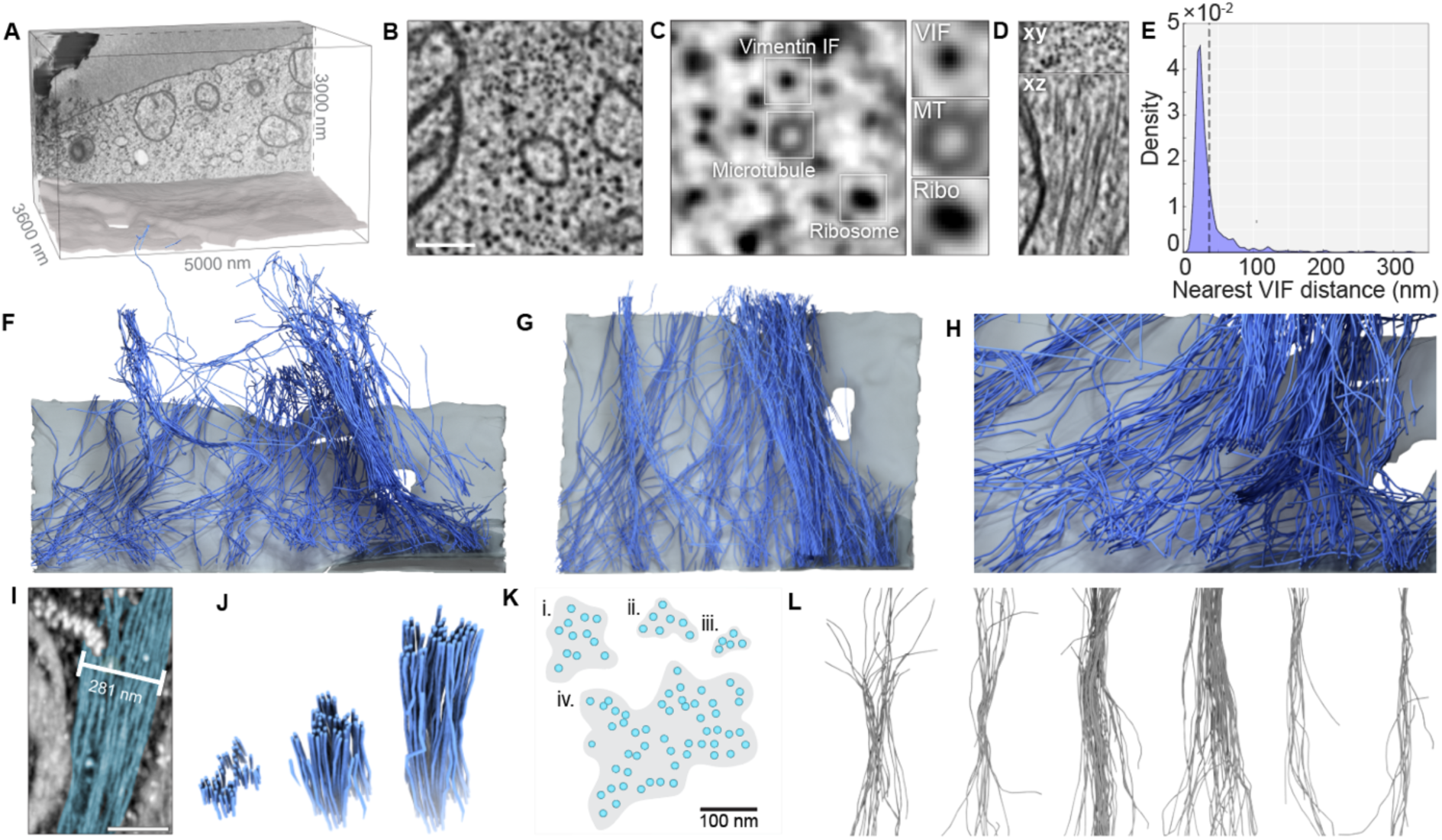
Reconstruction of vimentin filaments from FIB-SEM volumes reveals sparse, loosely organized vimentin bundles. **A.** 3D bounding box that corresponds to the analyzed FIB-SEM volume. A single XY slice from the rear of the volume is projected on the back wall and a surface rendering of the ventral plasma membrane is shown below. **B.** A single 800 x 800 nm FIB-SEM slice from this volume. Scale bar= 200 nm. **C.** Enlarged FIB-SEM slice highlighting the cross-sectional profile of a VIF (top), a microtubule (MT, center), and a ribosome (Ribo, bottom**). D.** Orthogonal views of vimentin short (top) and long (bottom) axes. **E.** Probability density estimate of vimentin-vimentin first nearest neighbor distances from each of 273,000 vertices derived from 583 VIF tracings. (**F-H**) Orthographic 3D renderings of vimentin (blue) and ventral plasma membrane (gray) within the volume indicated in a. **I.** Maximum intensity projection of a raw FIB-SEM stack featuring a vimentin bundle (blue overlay). Scale = 200 nm. **J.** 3D rendering of a vimentin bundle clipped at various distances along its length. **K.** Segmentations of four VIF bundle cross-sections derived from a single FIB-SEM slice. Blue dots indicate single filaments, shaded regions indicate cross-sectional area of bundles. **L.** Aligned montage of vimentin bundles derived from the volume indicated in A.

VIFs were identified in the FIB-SEM volume based on their cross-sectional short axis profile, where they appeared as uniform intensity ∼12 nm spots in the cytoplasm (Fig. 5 B and C; Fig S3B). VIF profiles most closely resembled ribosomes, though they measured smaller in width (11.9 nm VIF, 21.0 nm ribosome) and displayed slightly weaker staining intensity (Fig. 5 C; Fig. S3 C-D). VIFs were easily distinguishable from microtubules, based on the latter’s characteristic 24 nm annular cross-section which was readily resolvable in the 2 nm voxel EM dataset. Though we cannot exclude the possibility that these filaments represent copolymers between vimentin and other intermediate filaments such as nestin or peripherin, RNAseq indicates that vimentin is the dominant intermediate filament expressed in this cell type. When viewed orthogonally to their short axis, we observed elongated VIFs extending over thousands of slices, allowing us to track their centroids across, in many cases, the entire 3D region of interest (Fig. 5 D).

Within the 5 µm × 3 µm × 3.6 µm volume, we identified 583 VIFs representing over a millimeter of total filament length (Video 7). These VIFs closely interacted with one another and were broadly coaligned along the y axis. Computational analysis of VIF nearest neighbor distances revealed a mean inter-filament distance of 36 nm (Fig. 5 E) with a pronounced positive skew likely due to the presence of several isolated filaments in the remote corners of the volume. In 3D reconstructions, we observed that VIFs formed an intricate and locally co-oriented network characterized by overlapping and loosely interconnected bundles. Bundles varied widely in width, filament count, and cross-sectional filament density (Fig. 5 D-H). Disorganized, loosely aligned filaments coalesced to form ordered, lateral arrays of as many as 50 VIFs, before fraying out and diverging into multiple smaller bundles. Consistent with published in vitro data, vimentin filaments with bundles appeared stiffer and less curved than single unbundled filaments.

### VIFs are dynamically coupled to microtubules at intermittent crossover sites

Within the annotated perinuclear EM volume, we also identified and reconstructed 39 microtubules (Fig. 6 A-C). Given the co-alignment between vimentin and microtubule networks observed in light microscopy experiments (Gyoeva & Gelfand, 1991; Gan et al., 2016) we predicted that VIF bundles would be tightly associated – potentially even directly scaffolded – by microtubules. To our surprise, we observed that the two networks did not substantially co-distribute at the nanoscale.

**Figure 6:**
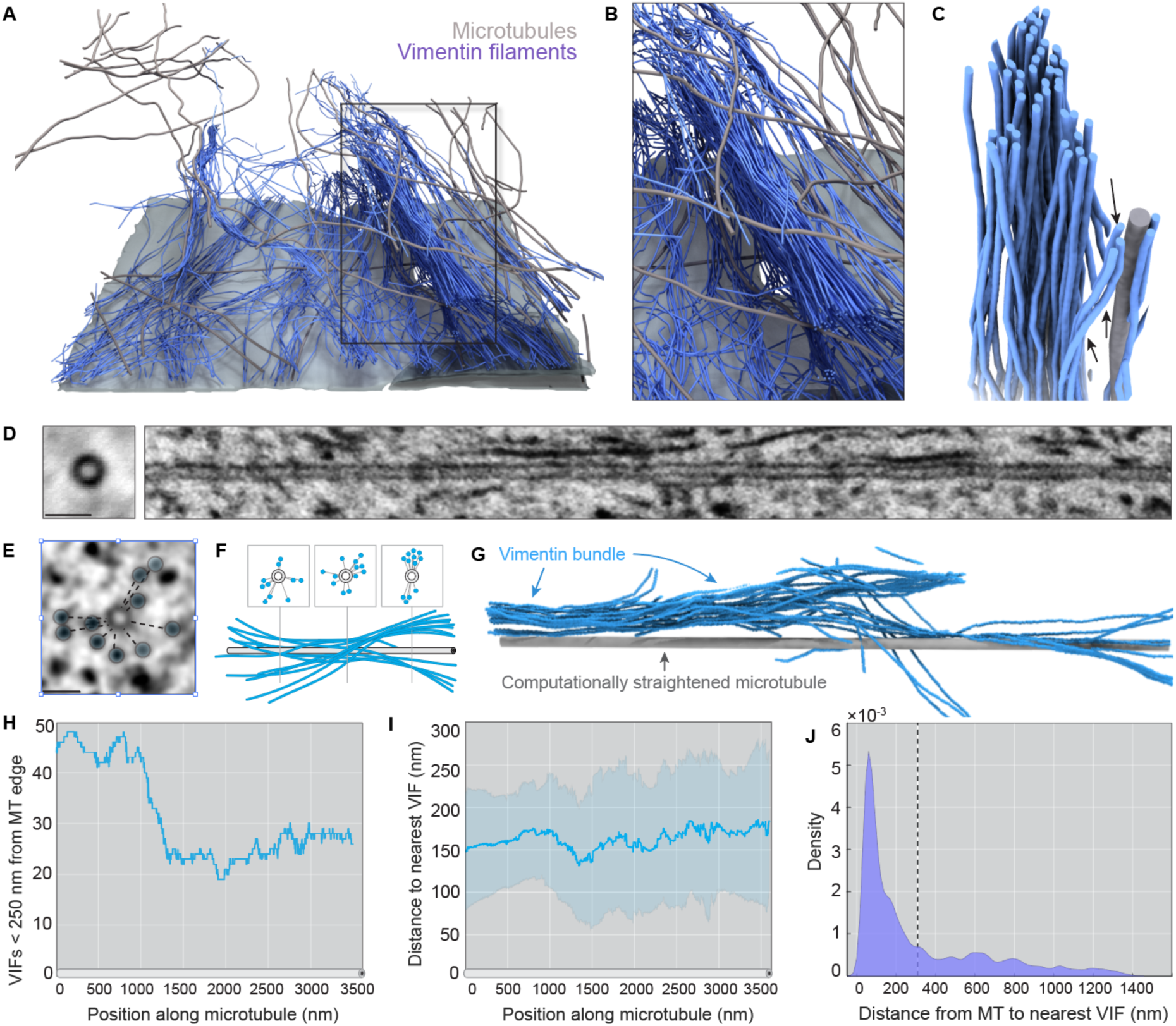
Vimentin filaments are loosely coupled to microtubules. a. 3D rendering of vimentin filaments (blue) with microtubules (gray) and ventral plasma membrane (light gray) in the target volume specified in Fig 5a. b. Enlarged view of a vimentin bundle (blue) and surrounding microtubules (gray). c. 3D reconstruction of vimentin filaments (blue) and a neighboring microtubule (gray). Filaments within the core of the bundle (black arrows) are positioned over 100 nm from the surface of the microtubule. Several filaments appear to diverge from the central bundle in order to align with the microtubule (black arrowheads). d. (left) Sum intensity projection of a center aligned 3500 nm microtubule trajectory. Scale bar = 50 nm. (right) Orthogonal view of MT center aligned track. By aligning the stack around the microtubule centroid, we have computationally straightened the microtubule, allowing for visualization of the local neighborhood around a single microtubule. e. Microtubule centered FIB-SEM slice with VIF cross-sections highlighted by blue circles and distances from the microtubule delineated by the dashed lines Scale bar = 50 nm. f. Cartoon schematic relating the XY view of microtubules and vimentin as seen in e to the expected appearance in the orthogonal view. g. 3D reconstruction of a computationally straightened microtubule (gray) and local vimentin filament bundle (blue). h. Total number of vimentin filaments within 249 nm of the centroid of a microtubule over its 3500 nm length. (Note: 249 nm from the center corresponds to 235 nm from the edge as MTs have a radius of 12 nm). i. Average distance of vimentin filaments from a single microtubule over its 3500 nm length within the reconstruction volume. Line = mean, error bars = standard deviation. j. Probability density estimates of distances between ∼30,000 MT vertices and their nearest vimentin neighbors. Line indicates a mean distance of 309 nm.

Three-dimensional analysis of the relative position and orientation of all vimentin and microtubule nearest neighbor vertices revealed limited spatial proximity or co-alignment between the two cytoskeletal networks. Using a computational straightening approach, we generated microtubule-centered sub-volumes, allowing us to visualize the organization and proximity of VIFs relative to the microtubule edge (Fig. 6 E-G and Video 7). Nearest neighbor analysis of ∼30,000 microtubule vertices revealed an average distance of 309 nm to the closest vimentin surface (Fig. 5 H-I; standard deviation = 324 nm). Indeed, over 85% of all microtubule voxels were positioned at greater than 50 nm from the nearest VIF and we did not observe any examples in which a microtubule appeared to scaffold or support the core of a vimentin bundle. Analysis of VIF-to-microtubule nearest neighbor distance revealed a similar mean of 291 nm (S.D. = 213 nm). Thus, VIFs and microtubules do not form dense, interpenetrating networks as have been described for vimentin and actin filaments (Wu et al., 2022).

While this limited nanoscale VIF/microtubule spatial proximity seemed at first inconsistent with the clear macroscale relationship between vimentin and microtubule cytoskeletons (Gan et al., 2016), we reasoned that direct and sustained physical interaction between fibers is not necessarily required for cytoskeletal crosstalk. In theory, only a small interaction surface relative to the lengths of either filament would be sufficient to coordinate global alignment of VIFs and microtubules. Indeed, recent in vitro reconstitution work examining microtubule-vimentin interactions concluded that VIFs display transient, highly dynamic coupling to microtubules with only limited co-alignment (Schaedel et al., 2021). Furthermore, vimentin may never enter close proximity with microtubules if the two are joined by an extended linker such as large molecular motor or an extended cytolinker such as plectin. Thus, we focused our attention on potential transient VIF-MT cross-over sites, and asked whether we could observe a structural signature that might indicate VIF-microtubule pairing.

At VIM-MT crossovers, we noted that the majority of VIFs passed by the microtubules without any indication of direct interaction. However, a small number of VIFs display high curvature at these sites, bending away from previously aligned VIF neighbors, and locally realigning with the stiff microtubule (see Fig. 6 C and Video 8). We reasoned that the broad coupling between vimentin and microtubule cytoskeletons may be partially explained through these low frequency crossover events between single VIFs emanating from bundles. These points of contact, though infrequent when measured against the total filament surface area, may be sufficient to synchronize VIFs and MTs.

## Discussion

In this study, we employed the multimeric fluorescent SunTag labeling approach to sparsely tag and identify single vimentin filaments in live cells. We confirmed vimentin-SunTag integration into single VIFs using SIM imaging and platinum replica electron microscopy and validated that its expression did not disrupt the organization of the endogenous VIF cytoskeleton. Using this reporter, we tracked single VIFs in all regions of the cytoplasm, including in the previously inaccessible, dense packed perinuclear vimentin network. Surprisingly, even in these extraordinarily dense regions, vimentin-SunTag showed extensive motility, a level of VIF dynamics not previously observed or appreciated with other imaging techniques. Vimentin-SunTag imaging enables the monitoring of both global and localized VIF dynamics in real-time, facilitating the precise study of VIF dynamics in various cellular contexts, both globally and at specific regions of interest.

We consistently noted multiple vimentin-SunTag spots within the same bundle exhibiting independent movement, occasionally even in opposite directions. Concurrently, some particles within the same bundle remained either stationary or underwent short displacements. These observations represent the first evidence that the motility of vimentin filaments within a given bundle is often uncoordinated and suggest that the lateral attractive forces holding the bundled filaments together can be readily overcome by motor-based pulling. Supporting this weak VIF bundling model, 3D reconstructions of VIF bundles revealed loose-packed arrays with fluctuating filament number and packing density. VIF bundles appeared frayed, unraveling along their length as single VIFs intermittently peeled away from the bundle periphery to transiently realign with neighboring microtubules. Of note, these vimentin-microtubule junctures were brief and infrequent compared to the total length of each filament. In general, we observed a marked separation between bundled VIFs and microtubules with surprisingly low levels of direct interaction. The sporadic nature of this physical vimentin-microtubule coupling may account for how motors and their cargoes can successfully traverse seemingly impenetrably dense regions of the vimentin filament cytoskeleton. In light microscopy movies the subtle offset between VIF bundles and underlying microtubules is not as clearly apparent as in our 3D reconstructions from FIB-SEM volumes. Were the VIF bundles to instead enmesh microtubule tracks, this tight coupling could potentially occlude molecular motor binding and stall organelle trafficking leading to defects in subcellular organization. Together, our novel approaches to visualize vimentin reveal that VIF bundles are not tightly crosslinked structures akin to dense actin bundles, but rather weakly associated filament arrays from which single VIFs can freely enter and exit.

Based on our 3D reconstruction and vimentin-SunTag tracking, we can envision several mechanisms by which VIFs could either dampen or stimulate intracellular trafficking in response to cellular needs. On one hand, a dense intracellular vimentin mesh may act as a brake or anchor on microtubule-based organelle motion. In this scenario, VIFs may interact with passing organelles, either directly via a molecular linker or indirectly through nonspecific obstruction, slowing or stalling organelle motion via increased drag or occlusion of motor binding sites. Alternatively, given the binding affinity of vimentin for both motor proteins (Gyoeva and Gelfand, 1991; Robert et al., 2019; Helfand et al., 2002; Prahlad et al., 1998) and membranous organelles (Vitali et al., 2023; Nekrasova et al., 2011; Biskou et al., 2019; Schwarz and Leube, 2016), it is not implausible that vimentin filaments could also behave as motor adaptors. By providing multiple binding sites along its length, a vimentin filament could parallelize the engagement of motor proteins and thereby stimulate robust directed transport of a given cargo. Further, the regular, repeating geometry of a vimentin filament may also serve to array multiple cargoes or scaffold motor teams onto structures that would otherwise lack sufficient surface area to accommodate such bulk. Lastly, VIFs may even accommodate motor-based sliding of single or multiple microtubules and therefore play important roles in cytoskeletal patterning. This concept, that vimentin may act not simply as passive cargo, but as a dynamic versatile motor adaptor, opens exciting new possibilities towards understanding the complex regulatory mechanisms underlying intracellular trafficking.

Cellular transport is the coordinated function of both microtubule and actin-based systems (Langford, 1995; Rogers and Gelfand, 1998). The long linear transport of vimentin-SunTag-labeled single filaments is lost after depolymerization of microtubules. Additionally, the disruption of microtubule motor function through either genetic modification or chemical inhibition leads to a significant decrease in long-range anterograde and retrograde transport. These observations suggests that transport of VIFs by the plus-end microtubule motor kinesin-1 and the minus-end motor cytoplasmic dynein are coordinated, a property that we observed in the past for the transport of membrane organelles (Kural et al., 2005; Hendricks et al., 2010; Barlan et al., 2013).

Previous studies have extensively documented interactions between VIFs and actin filaments (Hollenbeck et al., 1989; Wu et al., 2022; Esue et al., 2006). The possibility arises that the short-range movement of VIFs may depend on the actin cytoskeleton and its associated motor proteins. Vimentin-SunTag can serve as an excellent tool to address and investigate this intriguing question, providing a platform to explore the interplay between VIFs and the actin cytoskeleton in cellular dynamics. We hypothesize that the occurrence of short pauses and directional reversals in VIF/cargo transport along microtubule tracks may be the result of a molecular tug-of-war or a coordinated action between microtubule motors (kinesin-1 and dynein) and actin motors (myosin) (Evans et al., 2014; D’Souza et al., 2022; Vale, 2003). The mechanisms underlying this molecular tug-of-war are not fully understood; however, we believe that specific adaptors or linker proteins bind to cargoes.

## Materials and methods

### Plasmids

The SunTag system plasmids pcDNA4TO-K560-E236A-24xGCN4_v1-IRES-Puro (Addgene plasmid #60909; RRID: Addgene_60909) and pHR-scFv-GCN4-sfGFP-GB1-NLS-dWPRE (Addgene plasmid #60906; RRID: Addgene_60906) were generously provided by Ron Vale (Tanenbaum et al., 2014). The K560-E236A sequence in pcDNA4TO was replaced with vimentin cDNA tagged with a 3X flag tag, resulting in the fusion of a 24xGCN4 peptide epitope to the C-terminus of vimentin-3X flag sequence. To achieve low expression levels of vimentin fused with the 24xGCN4 peptide, this construct was sub-cloned into the pLVX-CMV100 lentiviral plasmid (truncated CMV promoter), which was kindly provided by Kevin Dean and Reto Fiolka (Addgene plasmid #110718; RRID: Addgene_110718) (Dean et al., 2016). The use of a truncated CMV100 promoter allowed for controlled expression of vimentin fused with 24xGCN4 peptide repeats, facilitating sparse SunTag labeling. In the pEGFP-1 plasmid backbone, the human dynamitin (p50) cDNA was inserted between the EcoRI and BamHI sites, and the EGFP cDNA was replaced with the mCherry cDNA. A vimentin-tagged mCherry plasmid was generated by replacing the p50 with vimentin cDNA in the same plasmid between the EcoRI and BamHI sites.

### Cell cultures

Human retinal pigment epithelial (RPE) cells were cultured in DMEM (Sigma, #D5648) supplemented with 10% fetal bovine serum and maintained at 37°C with 5% CO_2_. The kinesin-1 (KIF5B) knockout cell line was generated in our lab earlier using the CRISPR technique (Robert et al., 2019). For imaging experiments, cells were seeded onto clean glass coverslips in 35 mm dishes. The following day, plasmid transfections were performed using Lipofectamine™ 3000 transfection reagent (Thermo Fisher, #3000-008) or ViaFect (Promega, E498A) following the manufacturer’s protocol. Imaging was conducted 24 hours post-transfection. To induce microtubule depolymerization, cells were treated with 10 µM Nocodazole. Cytoplasmic dynein was inhibited using 5 µM Dynapyrazole A (Tocris, #6896).

### Live cell imaging details

Live cell time-lapse sequences were captured using a Nikon Eclipse Ti2 stand equipped with a W1 spinning disk confocal head (Yokogawa CSU with a pinhole size of 50 μm), utilizing either a 60× 1.49 N.A. or a 100× 1.45 N.A. oil immersion lens. Image acquisition was performed with a Hamamatsu ORCA-Fusion Digital CMOS Camera (Model: C13440-20CU) driven by Nikon NIS-Elements software (Version 5.42.01). A Tokai-Hit stage-top incubator and an Okolab gas mixer were employed to maintain a constant temperature of 37°C and a CO_2_ level of 5%.

### Immunostaining for structured illumination microscopy

After 24 hours of transfection, cells were pre-fixed for 3 minutes at 37 °C using a 0.3% glutaraldehyde and 0.2% Triton X-100 in PHEM buffer (60 mM PIPES, 27 mM HEPES, 10 mM EGTA, 8 mM MgSO_4_·7H_2_O, pH 7.0). Subsequently, cells were post-fixed for 15 minutes at 37 °C using 4% paraformaldehyde in PHEM buffer. Then free aldehyde groups were quenched with sodium borohydride (5 mg/ml) for 5 minutes, and cells were washed three times for 5 minutes each with PBS. Then cells were then incubated with primary antibody against vimentin (Chicken, Biolegend #919101, 1:2000) for 45 minutes at room temperature after blocking with 2.5% BSA. Subsequently, cells were incubated with secondary antibody Alexa-Fluor 647-conjugated donkey anti-chicken (Jackson Immuno Research #703-605-155, 1:700) for 45 minutes. Finally, coverslips were mounted and allowed to cure for 24 hours at room temperature before imaging.

Images were acquired using the CrestOptics DeepSIM-X-light (#S8002) mounted on the Nikon Eclipse Ti microscope stand, equipped with a Plan Apo λ 60× /1.4 N.A oil immersion objective lens. The Celesta light engine (Lumencor, #80-10268) served as the light source. Light emitted from the samples passed through specific bandpass filters (Chroma, blue channel: #392685, green channel: #417501, red channel: #414379, far-red channel: #417418) corresponding to their respective fluorescent channels. Images were captured by a Teledyne photometric BSI sCMOS camera controlled by Nikon NIS-Elements software (Version 5.42.03). In the DeepSIM module (65 images per focal plane), Z-stack images were acquired at intervals of 0.1 µm/step. The raw images were reconstructed into SIM images using the SIM reconstruction module in Nikon NIS-Elements.

### STED microscopy

COS7 cells plated on #1.5 glass bottom dishes (MatTek) were fixed in –20 °C methanol for 5 minutes and washed 3x in PBS. Cells were then incubated in PBS with 2% BSA containing primary antibodies against vimentin (Chicken, Encor CPCA-Vim, 1:1000) and alpha-tubulin (Mouse, Sigma T6199, 1:1000) overnight at 4 °C. Cells were washed 3x in PBS and incubated with goat anti-mouse AlexaFluor 488-plus (Invitrogen, A28175, 1:1000) and goat anti-chicken AlexaFluor 555-plus (Invitrogen, A32932, 1:1000) for 45 minutes. Samples were imaged on a Leica SP8 STED microscope using a 100x/1.4 NA objective lens. Samples were excited with a tunable white light laser set to 488 or 555 nm and STED was performed using frame scanning with first 660nm then 592nm depletion lasers. Raw images were processed in Fiji using a 1.5-pixel gaussian blur followed by contrast limited adaptive histogram equalization with block size set to 64 pixels and maximum contrast set to 2.0.

### Immunogold staining and electron microscopy of platinum replica

Cells labeled with vimentin-SunTag were extracted and fixed following the previously described procedure (Renganathan et al., 2023). In detail, cells were grown overnight on glass coverslips. The membrane and soluble cytosolic proteins were extracted with 1% Triton-X100 in PEM buffer (80 mM Pipes, 1 mM EGTA, 1 mM MgCl_2_ at pH 6.8) for 5 min. Cells were treated with 0.6 M potassium chloride in PHEM buffer, for 10 min to remove microtubule and actin filaments from the cytoskeletons still attached to the coverslips. Samples were fixed with 2% glutaraldehyde in 0.1 M cacodylate buffer. Subsequently, the cells were incubated with a rabbit anti-GFP antibody (Abcam, #6556) and then probed with a secondary antibody tagged with 12 nm colloidal gold (Jackson Immunoresearch, #111-205-144). Platinum replica electron microscopy (PREM) was performed as previously described (Svitkina, 2009; Korobova and Svitkina, 2008). Samples were imaged using a FEI Tecnai Spirit G2 transmission electron microscope (FEI Company, Hillsboro, OR) operated at 80 kV. Images were captured by Eagle 4k HR 200kV CCD camera and presented in original contrast.

### Image processing and analysis

The images were initially processed using ThunderSTORM, an image processing tool for particle localization (Ovesný et al., 2014). To determine the density of vimentin-SunTag particles per unit area, we applied a threshold of 2 times the standard deviation (2*std) (Wave.F1) to identify local maxima. Subsequently, we selected a region of interest (ROI) measuring 110 pixels by 110 pixels in three distinct areas of the cell to quantify the number of spots within each ROI. This ROI corresponds to an area of 30.25 μm². The number of spots in the ROI was divided by its area to estimate spot density per square micrometer.

For tracking vimentin-SunTag particles, we utilized the ThunderSTORM plugin, applying wavelet filtering (B-Spline, scale 2.0, order 3) and local maximum detection (8-neighbourhood, threshold at 1 time the standard deviation (1*std) for particle identification. The script then performed particle tracking using a total squared displacement minimization subroutine (Crocker and Grier, 1996), calculating various parameters such as maximum displacement, total distance traveled, velocity, and mean speed for each tracked particle.

### FIB-SEM Sample Preparation, Embedding and Imaging

COS-7 cells (ATCC; CRL-1651) were cultured on sapphire coverslips and subjected to high-pressure freezing (HPF) using a Wohlwend HPF Compact 02 machine. HPF was immediately followed by freeze substitution in a media containing 2% osmium tetroxide, 0.1% uranyl acetate, and 3% water in acetone, performed in an automated freeze substitution system (AFS2, Leica Microsystems) with a controlled temperature ramp from −140°C to room temperature over 39 hours. Subsequently, the samples were embedded in Durcupan ACM resin following a series of dehydration and infiltration steps and polymerized at 65°C for 48 hours.

For FIB-SEM imaging, resin-embedded samples were trimmed to reveal the region of interest and sputter-coated with a dual layer of 10 nm gold followed by 100 nm carbon to enhance conductivity. Imaging was performed on a custom Zeiss FIB-SEM system equipped with a Zeiss Capella FIB column and a Gemini 500 SEM. The block face was sequentially imaged and milled using a focused gallium ion beam. The imaging was performed with a 250-pA electron beam with a landing energy of 0.9 keV and a pixel size of 2 nm, while the milling was conducted with a 15 nA gallium ion beam removing 2 nm of material per cycle. This process was repeated continuously over seven days to acquire a near-isotropic volumetric dataset.

### FIB-SEM Post-Processing

Following acquisition, the raw FIB-SEM image data underwent -processing steps including field flattening to compensate for non-uniform detector sensitivity and signal fusion to integrate data from secondary electron and back-scattered electron detectors. Image registration and alignment were performed using a Scale Invariant Feature Transform (SIFT) algorithm with a Regularized Affine Transformation model to ensure accurate reconstruction of the three-dimensional volume.

### 3D Reconstruction of Filaments

Vimentin and microtubule trajectories were manually traced on 2 x 2 x 2 nm voxel FIB-SEM volumes in KNOSSOS. Using a custom python script, XYZ trajectories for each of 583 vimentin filaments and 39 microtubules were extracted, linearly interpolated with uniform 4 nm sampling, and smoothed using Savitzky-Golay filtering with a 3^rd^ degree polynomial and window sizes of 23 points and 53 points for vimentin and microtubules, respectively. For visualization, the coordinates of smoothed, interpolated trajectories were written to a JSON file and imported into Blender 4.0 using a custom python script. In Blender, filaments were rendered as curve objects with bevel depth equivalent to the filament radii as measured in the raw FIB-SEM dataset (6 nm for vimentin, 12 nm for microtubules). For Movie 7, a custom python script was used to randomly pseudocolor the VIFs. All renders were generated using the cycles render engine.

## Statements and Declarations

All authors declare no conflict of interest.

## Acknowledgements and Funding Information

The authors thank Dr. Stephen A Adam (Northwestern University) for stimulating discussions and critical reading of the manuscript and Dr. Farida Korobova, Center for Advanced Microscopy (Northwestern University) for PREM sample preparation and EM image acquisition.

The Gelfand lab is supported by the grant R35-GM131752 from the National Institute of General Medical Sciences.

The Jennifer Lippincott-Schwartz lab is supported by the Howard Hughes Medical Institute Janelia Research Campus.

Wei-Hong Yeo thanks the Christina Enroth-Cugell and David Cugell Fellowship for Visual Neuroscience and Biomedical Engineering at Northwestern University for their support.

Hao F. Zhang acknowledges National Institutes of Health grants R01GM139151, R01GM140478, and U54CA268084.

## Data Availability Statement

All data reported in this paper will be shared upon reasonable request.

---

**Supplementary Figure 1:**
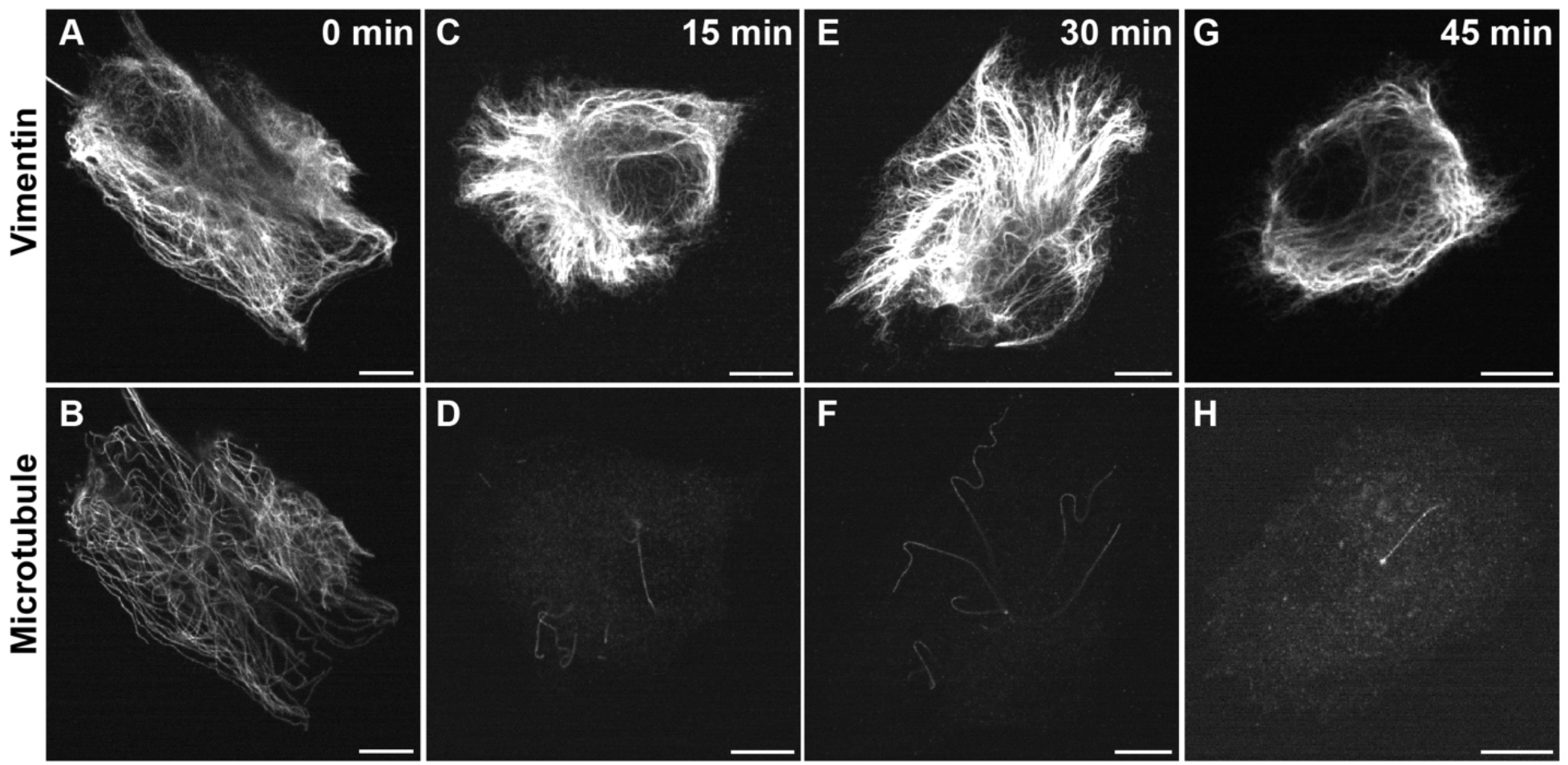
Vimentin briefly sustains its organization in the absence of microtubules. RPE cells were treated with 10 μM nocodazole for various time points. Subsequently, the cells were fixed using ice-cold methanol and subjected to immunofluorescence staining for vimentin and microtubules. Upper panel: Depicts the organization of vimentin at different time points following nocodazole treatment. Bottom panel: Displays the corresponding microtubule staining for the respective cells. Scale bar: 10 μm

**Supplementary Figure 2:**
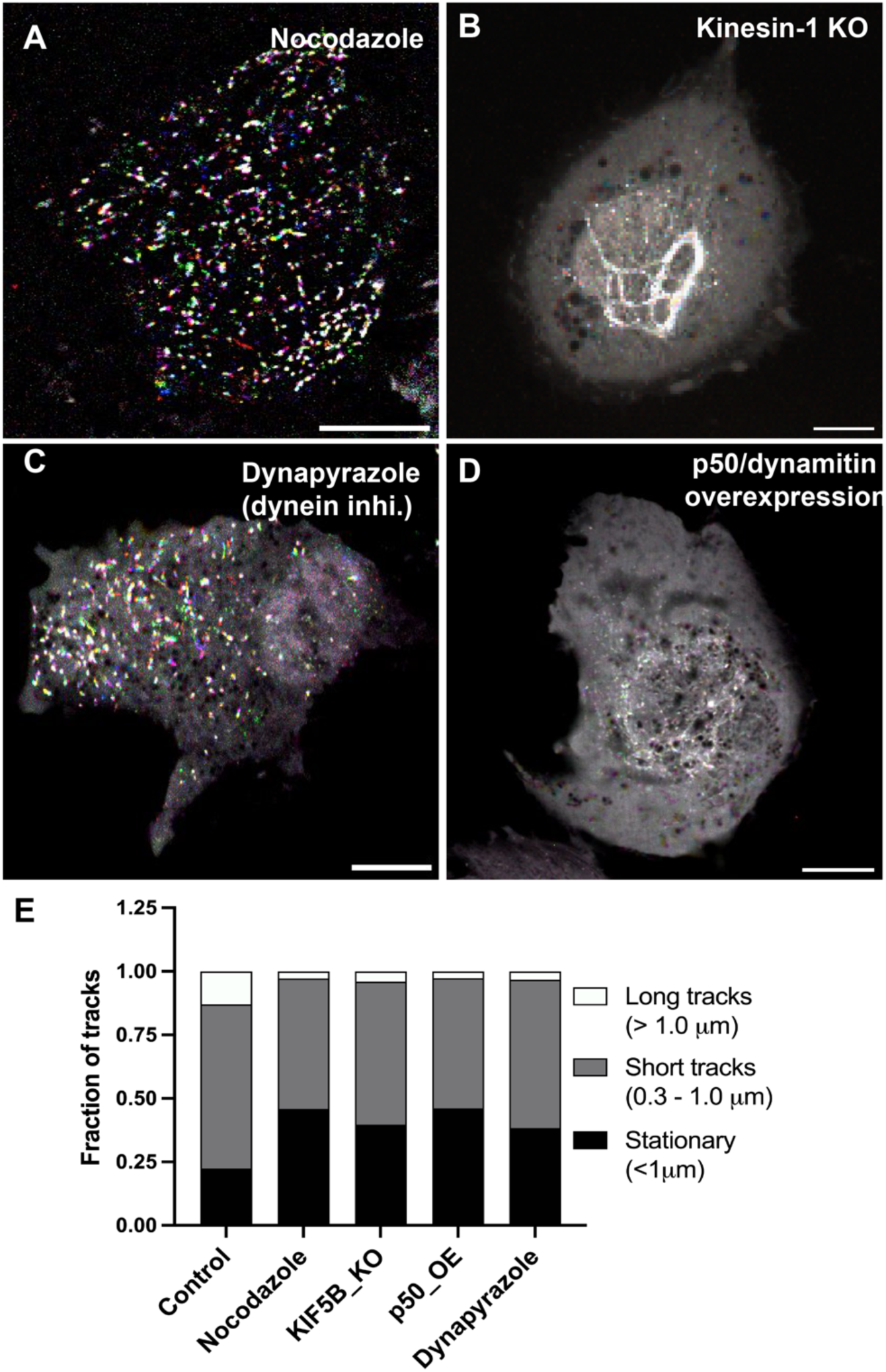
Vimentin filament motility along microtubule driven by microtubule motors kinesin-1 and cytoplasmic dynein. Vimentin-SunTag was labelled RPE cells, time-lapse sequence was collected for a total time of 1 minute at 2 fps and temporal color code generated respective conditions. Scale bar: 10 μm (A) Depolymerization of microtubule by 10 μM Nocodazole inhibits the vimentin-SunTag movement (see also Video S2). (B) In RPE kinesin-1 knock out (KO) cells, transport of vimentin-SunTag completely inhibited and aggregated to the perinuclear region (see also Video S3). (C) Inhibition of cytoplasmic dynein by 5 μM Dynapyrazole A blocks the vimentin-SunTag transport (see also video 4). (D) Overexpression of p50/dynamitin results in inhibition of vimentin-SunTag transport and aggregation in the perinuclear region like the KIF5B KO condition (see also Video S5). Scale bar: 10 μm. (E) Quantification of vimentin-SunTag motility categorized vimentin-SunTag motility based on the total displacement in microns among various conditions with. (*n* = 20 individual cells).

**Supplementary Figure 3:**
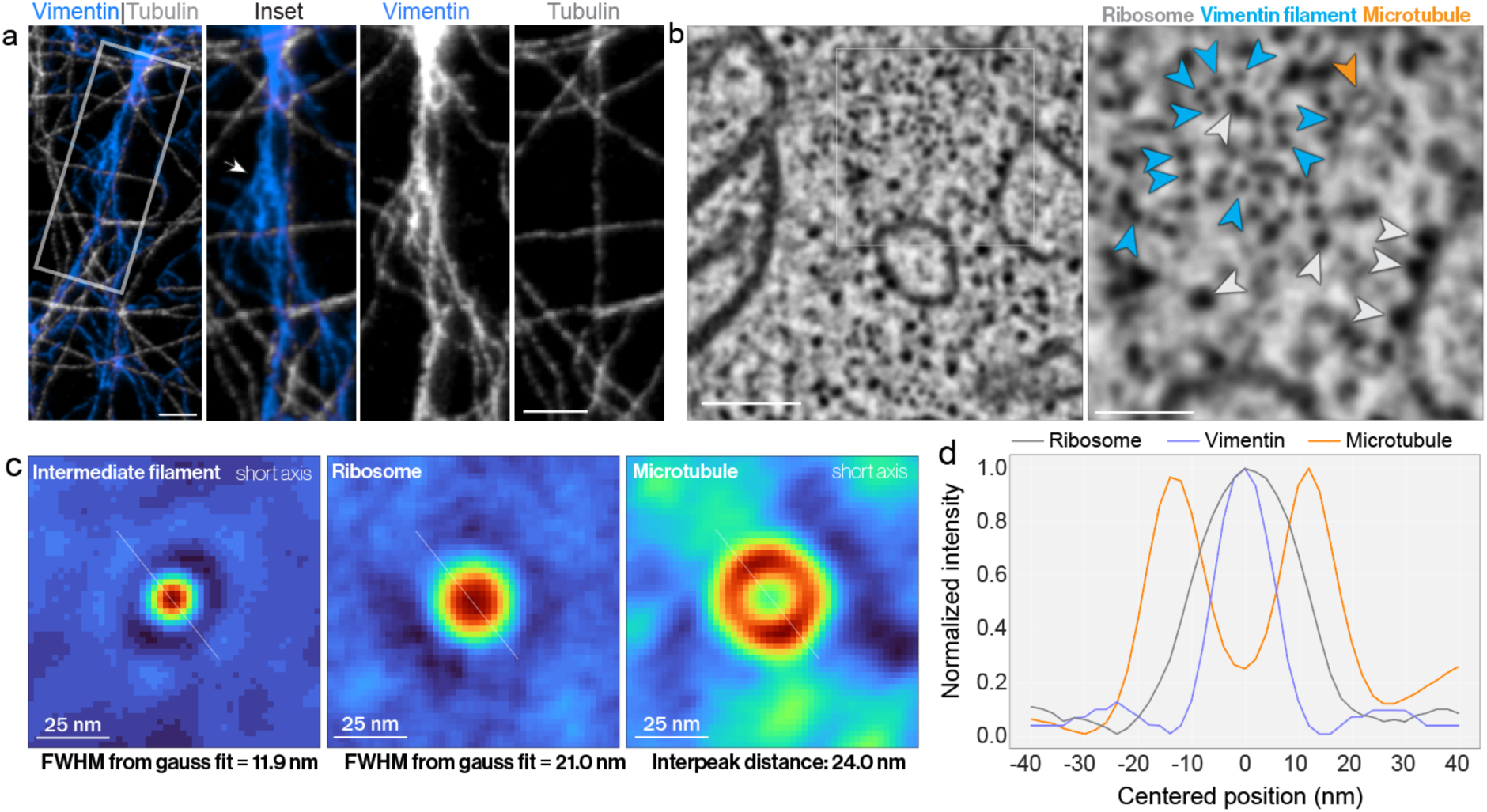
Relationship between vimentin and microtubules is ambiguous. **A)** Stimulated Emission Depletion (STED) microscopy image of immunostained vimentin intermediate filaments (blue) and microtubules (white) in a COS-7 cell. At the ∼ 50 nm measured resolution of this image, the organization of the vimentin bundle is obscured and the nanoscale relationship between vimentin and microtubules is ambiguous. **B)** FIB-SEM slice from Fig. 5B with inset indicating positions of vimentin intermediate filaments (light blue arrow), a microtubule (orange arrow), and ribosomes (light gray arrows). **C)** Center aligned average projections of VIFs (left), ribosomes (center), and microtubules (right) with measured size of each averaged structure indicated below. White diagonal lines indicate the position of line profile used in d. **D)** Intensity profile of averaged ribosome, vimentin, and microtubule profiles shown in c. Plots indicate mean intensity along each profile at 360 1-degree rotations.

## Video Captions

**Video 1**: Active Transport of vimentin-SunTag in Control Cell (RPE wild type): Suntag plasmids were co-transfected with mCherry-vimentin. The dynamic movement of active vimentin-SunTag (green dots) reflects the dynamics of vimentin filaments. The white box region is zoomed in, and the full-time sequence is subsequently displayed. vimentin-SunTag dots exhibit distinct phases, including long linear transport, short transit, and stationary vimentin-SunTag. Magenta color represents mCherry-vimentin.

**Video 2:** Active transport of vimentin-SunTag in control cell active vimentin-SunTag (green dots) movements represent the dynamics of vimentin filaments. A maximum time projection generated from the time sequence is displayed in the last 3 seconds of the video.

**Video 3:** Microtubule depolymerization inhibits the transport of vimentin-SunTag in RPE cells after treatment with 10 µm of Nocodazole, most vimentin-SunTag particles become stationary, showing inhibited transport. A maximum time projection generated from the time sequence is displayed in the last 3 seconds of the video

**Video 4:** Microtubule motor kinesin-1 loss inhibits the transport of vimentin-SunTag. After kinesin-1 KO, active transport of vimentin-SunTag is lost, and most particles cluster near the nucleus. There are no long linear trajectories, and most particles display confined movement. A maximum time projection generated from the time sequence is displayed in the last 3 seconds of the video

**Video 5:** Inhibition of cytoplasmic dynein by 5 µm of Dynapyrazole A causes loss of vimentin-SunTag transport. A maximum time projection generated from the time sequence is displayed in the last 3 seconds of the video

**Video 6:** Inhibition of cytoplasmic dynein by p50 overexpression in RPE cells leads to inhibition of vimentin-SunTag transport, similar to kinesin-1 KO cells, vimentin-SunTag dots cluster near the nucleus. A maximum time projection generated from the time sequence is displayed in the last 3 seconds of the video

**Video 7:** Three-dimensional reconstruction of 583 VIFs identified and tracked from the perinuclear FIB-SEM volume indicated in Fig 5a. Filaments are assigned a random color and progressively revealed to more clearly reveal the VIF network organization.

**Video 8:** Three-dimensional reconstruction of the 583 VIFs (white) and 39 microtubules (blue) reconstructed from the FIB-SEM volume indicated in Fig 5a.

